# Facioscapulohumeral muscular dystrophy is associated with altered myoblast proteome dynamics

**DOI:** 10.1101/2022.12.14.520394

**Authors:** Yusuke Nishimura, Adam J. Bittel, Connor A. Stead, Yi-Wen Chen, Jatin G Burniston

## Abstract

Proteomic studies in facioscapulohumeral muscular dystrophy (FSHD) could offer new insight to disease mechanisms underpinned by post-transcriptional processes. We used stable isotope (deuterium oxide; D_2_O) labelling and peptide mass spectrometry to investigate the abundance and turnover rates of proteins in cultured muscle cells from 2 individuals affected by FSHD and their unaffected siblings (UASb). We measured the abundance of 4485 proteins and the turnover rate of 2324 proteins in each (*n* = 4) myoblast sample. FSHD myoblasts exhibited a greater abundance but slower turnover rate of subunits of mitochondrial respiratory complexes and mitochondrial ribosomal proteins, which may indicate an accumulation of ‘older’ less viable mitochondrial proteins in myoblasts from individuals affected by FSHD. Our results highlight the importance of post-transcriptional processes and protein turnover in FSHD pathology and provide a resource for the FSHD research community to explore this burgeoning aspect of FSHD.

## Introduction

Facioscapulohumeral muscular dystrophy (FSHD) is the third most common muscular dystrophy and has a prevalence of approximate 1:20,000. Currently, there is no effective treatment or cure for FSHD, which is marked by gradual loss of muscle mass and function and eventual loss of independence (1). Ectopic expression of the double homeobox 4 (*DUX4*) gene is the key molecular cause of the primary (95% of cases) disease phenotype FSHD1 (OMIM 158900) and less common FSHD2 (OMIM 158901) (2). DUX4 instigates widespread changes in muscle gene expression (3), including aberrant activation of other transcription factors. Several biological processes are known to be disrupted by DUX4 expression in muscle, including myogenic differentiation and cell cycle (4), oxidative stress sensitivity (5), DNA damage (6), Wnt/β-catenin signaling (7), metabolic stress and mitochondrial dysfunction (8), and p53-mediated apoptosis (9). However, the mechanisms that connect DUX4 expression to muscle toxicity are not yet fully understood despite extensive studies on gene regulation and transcriptional processes (10–13).

Proteomic studies in FSHD are currently scarce but have the potential to bring new insight to the role of post-transcriptional processes in FSHD, and help connect dysregulation of gene programmes with cellular abnormalities. However, Jagannathan et al. (14) reported a striking disconnection between changes in gene expression and changes in the abundance of the corresponding proteins when DUX4 is artificially expressed in immortalized MB135 human myoblasts. After induction of DUX4, there was a greater than 4-fold change in the abundance of 208 out of 4,005 proteins studied, but when proteomic data were aligned with earlier gene expression data (15), one-third of changes in protein abundance were not matched by a change in gene expression or the change in expression of the gene was diametrically opposite to the change in protein abundance (14). This disconnection between mRNA and protein responses to DUX4 expression could occur, for example, through (mis)- regulation of synthetic processes, degradative processes, or a combination of the two. Artificial expression of DUX4 results in a 50 % reduction in bulk protein synthesis measured by ^35^S-methionine incorporation in M135 myoblasts (14). However, it is not clear whether similar disruptions in protein turnover occur in patient-derived samples, and the measurement of the ‘bulk’ synthesis rate of protein mixtures (14) cannot distinguish individual protein responses. The aforementioned 50 % reduction in ^35^S-methionine incorporation could represent a blanket 50 % reduction in the synthesis of all proteins or complete inhibition in the synthesis of 50 % of proteins. In addition, DUX4 target-genes that were faithfully translated into functional proteins included several known or putative E3 ubiquitin ligases that could dysregulate protein degradation (14).

DUX4 overexpression in a myoblast cell line (14, 16) is an optimised research model and may not faithfully reflect FSHD pathophysiology, where DUX4 expression is transient and often localized to small numbers of myonuclei at a particular time. Other proteomic analyses (17, 18) have used muscle samples of individuals with FSHD, including non-clinically affected muscles to focus on the fundamental pathophysiological mechanisms of FSHD whilst minimizing the impact of dystrophic processes. In addition, myotubes (19) or myoblasts (20) and interstitial fluid samples (21) from individuals with FSHD and their unaffected siblings (UASb) have been studied using proteomic techniques. Herein, we performed proteomic analysis on previously generated and validated (22) immortalized myoblasts from matched pairs of individuals with FSHD and UASb. To the best of our knowledge, the present work represents the first proteomic analysis of immortalized FSHD and UASb cell lines reported in Homma et al. (22), which have been characterized across several previous studies (8, 10, 23–27).

To study protein turnover, myoblasts were cultured with the stable isotope, deuterium oxide (D_2_O), which labels amino acid precursors and enables the fraction of newly synthesised protein to be calculated from time-dependent differences in the mass isotopomer distribution of peptide mass spectra (28, 29). Using this proteomics approach, we surveyed the abundances and turnover rates of thousands of proteins in FSHD and UASb cell cultures, and discovered some proteins exhibit a discordance between differences in abundance and turnover rate.

## Experimental Procedures

### Cell Culture and Deuterium Oxide Labelling

Immortalized human myoblasts from individuals with FSHD and UASb were obtained from the Senator Paul D. Wellstone Muscular Dystrophy Cooperative Research Center for FSHD at the University of Massachusetts Chan Medical School (Worcester, MA, USA) and Dr Woodring E Wright at the University of Texas Southwestern Medical Center. The collection of muscle biopsies, isolation of myoblast and purification, and molecular characterization of FSHD and UASb cells was originally described by Homma et al. (22).

An overview of the study design is shown in **Fig. 1A** and characteristics of FSHD and UASb donors are presented in **Fig. 1B**. Consistent with a previous work by Pandey et al. (30), immortalized FSHD myoblasts were cultured in LHCN medium (4:1 DMEM:Medium 199 (ThermoFisher Scientific, Waltham, MA, USA) supplemented with 15% fetal bovine serum (FBS, Hyclone, South Logan, UT, USA), 0.03 mg/mL ZnSO4 (Sigma), 1.4 mg/mL Vitamin B12 (Sigma-Aldrich), 2.5 ng/mL hepatocyte growth factor (Chemicon International, Temecula, CA, USA), 10 ng/mL basic fibroblast growth factor (Millipore, Billerica, MA, USA), and 0.02 M HEPES (Life Technologies, Carlsbad, CA, USA) with dexamethasone (140 nmol/ml) to suppress DUX4 expression and facilitate the proliferation of FSHD cells. After attaining 80% confluency, cultures were switched to LHCN media without dexamethasone (LHCN -DEX) for three days to wean myoblasts from the effects of dexamethasone.

**Figure 1.**
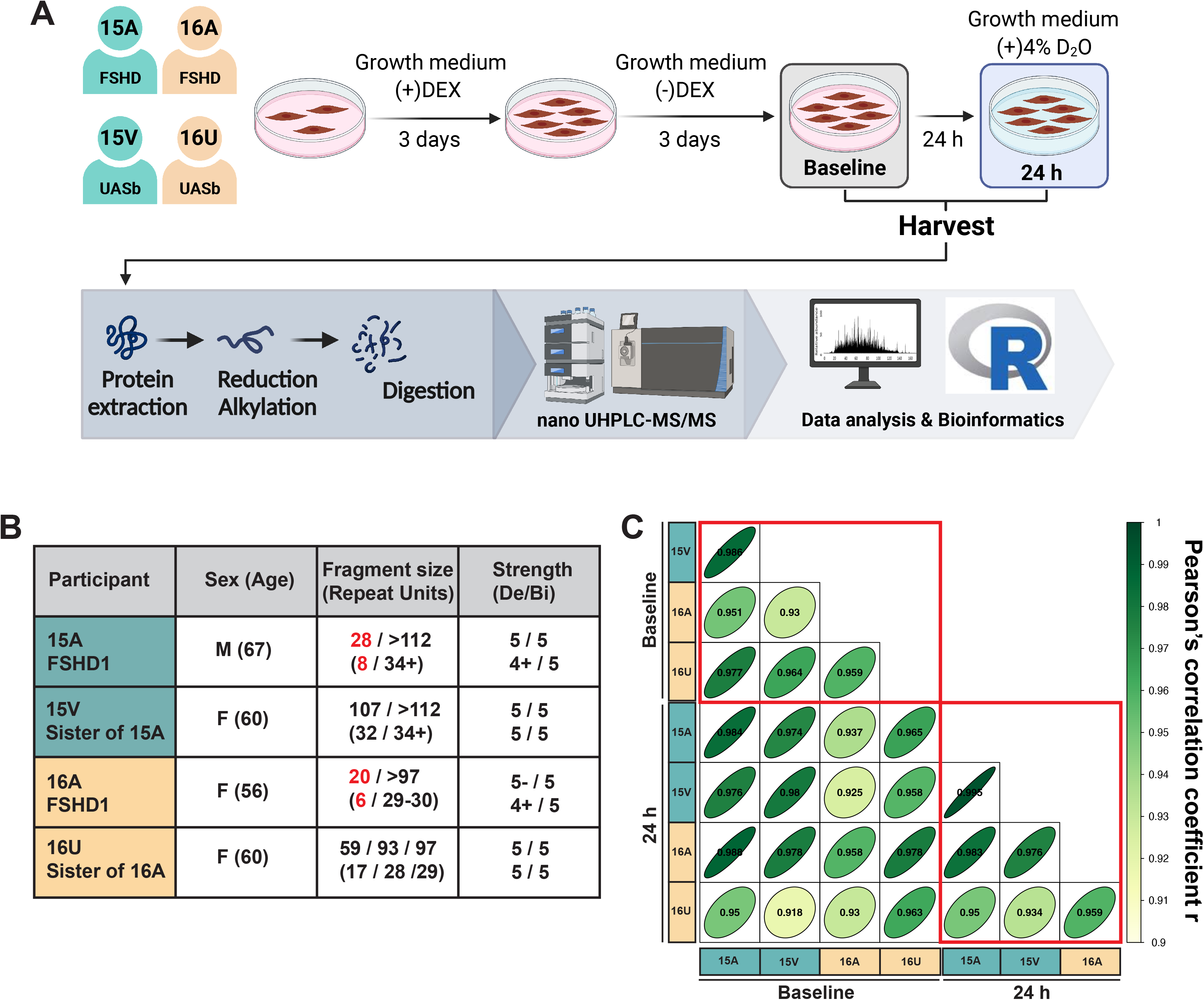
Dynamic proteome profiling of two matched pairs of FSHD and UASb myoblasts. **A)** Experimental design and workflow for sample preparation and analysis. **B)** Characteristics of FSHD and UASb donors, including shortened length of the 4q D4Z4 repeat array (bold red) and muscle score using the Medical Research Council (MRC) scale where 5/5 is maximum strength. Data was retrieved from Homma et al. (22). Del, deltoid; Bi, biceps. **C)** Matrix correlation in protein abundance of n = 4485 proteins. Pearson’s correlation coefficient was determined to identify the correlation in protein abundance.

Myoblasts were then cultured for an additional 24 hours in LHCN -DEX media supplemented with 4% v/v deuterium oxide (D_2_O) to label newly synthesized proteins, and then harvested for analysis. Myoblasts derived from UASb were treated identically to provide a control group at each experimental time point.

### Protein Extraction and Quantification

Following treatments and timings detailed above, medium was aspirated, and the cell monolayer was washed twice with ice cold PBS. Cells were lysed with 250 μl of Urea buffer (8 M Urea, 100 mM Tris, pH ~8.5) for 5 min at room temperature, scraped into Eppendorf tubes in preparation for total protein quantification, digestion, and proteomic analyses. Total protein concentration (μg/μl) was quantified against bovine serum albumin (BSA) standards using the Pierce™ BCA Protein Assay Kit (Rockford, IL, USA), according to the manufacturer’s instructions.

### Protein Digestion

Filter-Aided Sample Preparation (FASP) was performed using lysates containing 100 μg protein and incubated at 37 °C for 15 min in UA buffer with 100 mM dithiothreitol (DTT) followed by 20 min at 4 °C in UA buffer containing 50 mM iodoacetamide (protected from light). Samples were washed twice with 100 μl UA buffer and transferred to 50 mM ammonium hydrogen bicarbonate (Ambic). Sequencing grade trypsin (Promega; Madison, WI, USA) in 50 mM Ambic was added at an enzyme to protein ratio of 1:50 and the samples were digested overnight at 37 °C. Peptides were collected in 50 mM Ambic and trifluoracetic acid (TFA) was added to a final concentration of 0.2 % (v/v) to terminate digestion. Aliquots, containing 4 μg peptides, were desalted using C18 Zip-tips (Millipore, Billercia, MA, USA) and eluted in 50:50 of acetonitrile and 0.1 % TFA. Peptide solutions were dried by vacuum centrifugation for 25 min at 60 °C and peptides were resuspended in 0.1 % formic acid spiked with 10 fmol/ul yeast ADH1 (Waters Corp.) in preparation for LC-MS/MS analysis.

### Liquid Chromatography-Mass Spectrometry Analysis

Peptide mixtures were analysed using an Ultimate 3000 RSLC nano liquid chromatography system (Thermo Scientific) coupled to a Fusion mass spectrometer (Thermo Scientific). Samples were loaded on to the trapping column (Thermo Scientific, PepMap100, C18, 75 μm X 20 mm), using partial loop injection, for 7 minutes at a flow rate of 9 μl/min with 0.1 % (v/v) TFA. Samples were resolved on a 500 mm analytical column (Easy-Spray C18 75 μm, 2 μm column) using a gradient of 96.2 % A (0.1 % formic acid) 3.8 % B (79.9 % ACN, 20 % water, 0.1 % formic acid) to 50 % A 50 % B over 90 min at a flow rate of 300 nl/min. The data-dependent program used for data acquisition consisted of a 120,000-resolution fullscan MS scan (AGC set to 4e5 ions with a maximum fill time of 50 ms) with MS/MS using quadrupole ion selection with a 1.6 m/z window, HCD fragmentation with a normalized collision energy of 32 and LTQ analysis using the rapid scan setting and a maximum fill time of 35 msec. The machine was set to perform as many MS/MS scans as possible to maintain a cycle time of 0.6 sec. To avoid repeated selection of peptides for MS/MS, the program used a 60 s dynamic exclusion window.

### Label-Free Quantitation of Protein Abundances

Progenesis Quantitative Informatics for Proteomics (QI-P; Nonlinear Dynamics, Waters Corp., Newcastle, UK) was used for label-free quantitation, consistent with previous studies (31–34). Log-transformed MS data were normalized by inter-sample abundance ratio, and relative protein abundances were calculated using nonconflicting peptides only. In addition, abundance data were normalized to the 3 most abundant peptides of yeast ADH1 to obtain abundance estimates (ABD_mol_) in fmol/μg protein. MS/MS spectra were exported in Mascot generic format and searched against the Swiss-Prot database (2021_03) restricted to Homosapiens (20,371 sequences) using locally implemented Mascot server (v.2.2.03; www.matrixscience.com). The enzyme specificity was trypsin with 2 allowed missed cleavages, carbamidomethylation of cysteine (fixed modification), deamidation of asparagine and glutamine (variable modification) and oxidation of methionine (variable modification). M/Z error tolerances of 10 ppm for peptide ions and 0.6 Da for fragment ion spectra were used. The Mascot output (xml format), restricted to non-homologous protein identifications was recombined with MS profile data in Progenesis.

### Measurement of protein turnover rates

Protein fractional turnover rates (FTR) were calculated consistent with our previous work (33). Mass isotopomer abundance data were extracted from MS spectra using Progenesis QI (Nonlinear Dynamics, Waters Corp., Newcastle, UK). The abundance of m_0_–m_3_ mass isotopomers was collected over the entire chromatographic peak for nonconflicting peptides that were used for label-free quantitation. Mass isotopomer information was processed in R version 4.0.3 (R core team., 2016). Incorporation of deuterium into newly synthesized protein causes a decrease in the molar fraction of the monoisotopic (m_0_) peak.

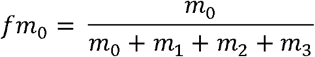

Equation 1: *fm*_0_ = molar fraction, *m*_0_ = monoisotopic peak, *m*_1_, *m*_2_, *m*_3_ = mass isotopomers 1 - 3.

Over the duration of the experiment, changes in mass isotopomer distribution follow a nonlinear exponential pattern as a result of the rise-to-plateau kinetics of D_2_O-labelled amino acids into newly synthesized protein. The rate constant (k) for the decay of fm_0_ was calculated as a first-order exponential spanning from the beginning (t) to end (?) of the 24 h labeling period, using Equation (2).

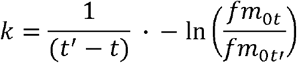

Equation 2: *k* = rate constant, *t* = first timepoint, *t′* = end timepoint, *fm*_0*t*_ = molar fraction at first timepoint, *fm*_0*t′*_ = molar fraction at last timepoint.

The rate of change in mass isotopomer distribution is also a function of the number of exchangeable H sites, and this was accounted for by referencing each peptide sequence against standard tables (35) that report the relative enrichment of amino acids by deuterium in humans. Peptide FTR was derived by dividing k by the molar percentage enrichment of ^2^H added to the culture media (p) and the total number (n) of ^2^H exchangeable H-C bonds in each peptide.

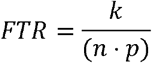

Equation 3: *k* = rate constant, *n* = number of H-D exchange sites, *p* = precursor enrichment

The median FTR of peptides assigned to each protein was used to calculate the FTR for each protein in each individual FSHD and UASb sample. Decimal values were multiplied by 100 to give FTR in %/h.

### Experimental Design and Statistical Rationale

The experiment was designed to investigate differences in the abundance and turnover rate of proteins that are common between two matched-pairs of FSHD and UASb control samples. All statistical analyses were performed using R version 4.2.1. Two-way mixed ANOVA was performed to assess protein abundances in FSHD (*n* = 2) and UASb (*n* = 2) samples at baseline (0h) and 24h time points and confirm that the abundance of most proteins was stable during the D_2_O labelling period. Subsequently, protein abundances measured at baseline and after 24 h of labelling with D_2_O were averaged for each biological replicate FSHD (*n* = 2) and UASb (*n* = 2) and one-way ANOVA was used to assess differences in protein abundance between FHSD and UASb groups.

Peptide mass isotopomer distributions measured at baseline and after 24 h of labelling with D_2_O were used to calculate protein turnover rates in each biological replicate FSHD (*n* = 2) and UASb (*n* = 2) and one-way ANOVA was used to assess differences in protein turnover rates between FHSD and UASb groups.

Differences in the abundance or turnover rate of proteins between FSHD and UASb groups are reported as log_2_ transformed data and statistical significance was set at *P* < 0.05. Due to the limited number of replicates (*n* = 2, per group) a false discovery rate criterion was not set, instead, q values (36) at the *P* = 0.05 threshold were reported.

### Bioinformatic Analysis

Gene ontology analysis of proteins more enriched (log_2_ Diff. >1) or depleted (log_2_ Diff. <-1) in FSHD was performed via Overrepresentation Enrichment Analysis (37) using the Gene Ontology enRIchment anaLysis and visuaLizAtion tool (Gorilla) (38, 39). Enrichment of GO terms was considered significant if the Benjamini and Hochberg adjusted *P*-value (40) was < 0.01. Protein interactions were investigated using bibliometric mining in the Search Tool for the Retrieval of INteracting Genes/proteins (STRING, Version 11.5) (41) and the interaction networks were illustrated using Cytoscape (Version 3.9.1) (42). The coverage of mitochondrial Complex subunit proteins and mitochondrial ribosomal proteins were surveyed as identified in Human MitoCarta 3.0. (43)

## Results

### Protein abundance profiling of FSHD and UASb myoblasts

Label-free quantitation encompassed 4485 proteins in myoblast cultures from two individuals affected by FSHD and their unaffected siblings (UASb), each sampled at 2 timepoints (baseline and after 24 hours labelling with D_2_O; **Fig. 1A**). Protein abundances were closely correlated (Pearson’s r ≥0.958) between sample pairs collected at baseline and 24 h timepoints, and correlations across FSHD and UASb samples, either within or between families, ranged between r = 0.93 and r = 0.995 (**Fig. 1C**). Two-way mixed analysis of variance between groups (FSHD vs UASb) and repeated over time (Baseline vs 24 h) found no statistical interactions (P values <0.05) that had a false discovery rate <69 %, and just 5 proteins (EEF2K, MRPL3, BRD3, HSPB3 and PODXL) with an interaction effect (P<0.05, q>0.69) exhibited a >2-fold difference (log_2_ Diff. −1< or 1>) in protein abundance. Eleven proteins (COX7A2L, PTX3, GINM1, MRPL48, UBR3, EFNB1, EEF2K, MRPL3, BRD3, PODXL, GJA1) exhibited a statistical main-effect of time (P values <0.05, q >0.48) and a >2-fold difference in protein abundance. Consequently, protein abundance was considered to be stable for the majority (99.4 %) of proteins during the 24 h D_2_O-labelling period, and in the analyses below the average of protein abundance values at baseline and 24 h timepoints were used.

One-way analysis of variance of time-averaged abundance data highlighted 114 proteins that exhibited a significant difference (P<0.05) between FSHD vs UASb groups, and 17 of these proteins exhibited a >2-fold difference in average protein abundance. Nine proteins (FKBP7, UFSP2, FHL1, EEF2K, USP11, PIP4K2C, NDUFA4, MRPL19, and UQCR11) were more abundant (log_2_ Diff. >1) in FSHD samples and 8 proteins (HAUS6, TBC1D22A, MAP3K7CL, FNBP4, CEP170B, CNOT9, TACC3, CCNA2) were less (log_2_ Diff. <-1) abundant in FSHD compared to UASb myoblasts (**Fig. 2A**). In addition, 6 proteins (DUSP23, ANAPC4, DDX56, PARP12, ZYG11B, MT-ND4) were detected exclusively in FSHD and 3 proteins (NPR3, CFB, FOXO1) exhibited extremely large (64-fold difference; log_2_ Diff. >6) abundance in FSHD compared to UASb. Twelve proteins (PHLDA1, ANKRD10, XAF1, DCAF1, DCLRE1C, IGFBP3, PRORP, GNL2, SLC25A23, SECTM1, TOP3B, FBP1) were uniquely detected in UASb samples and 3 proteins (ARNTL2, TRIL, MCM9) exhibited extremely low (−64-fold difference; log_2_ Diff. <-6) abundance in FSHD compared to UASb samples. Proteins that exhibited extremely large or low abundance differences between FSHD and UASb were due to family-specific differences between FSHD and UASb samples (**Supplemental Fig. S1**).

**Figure 2.**
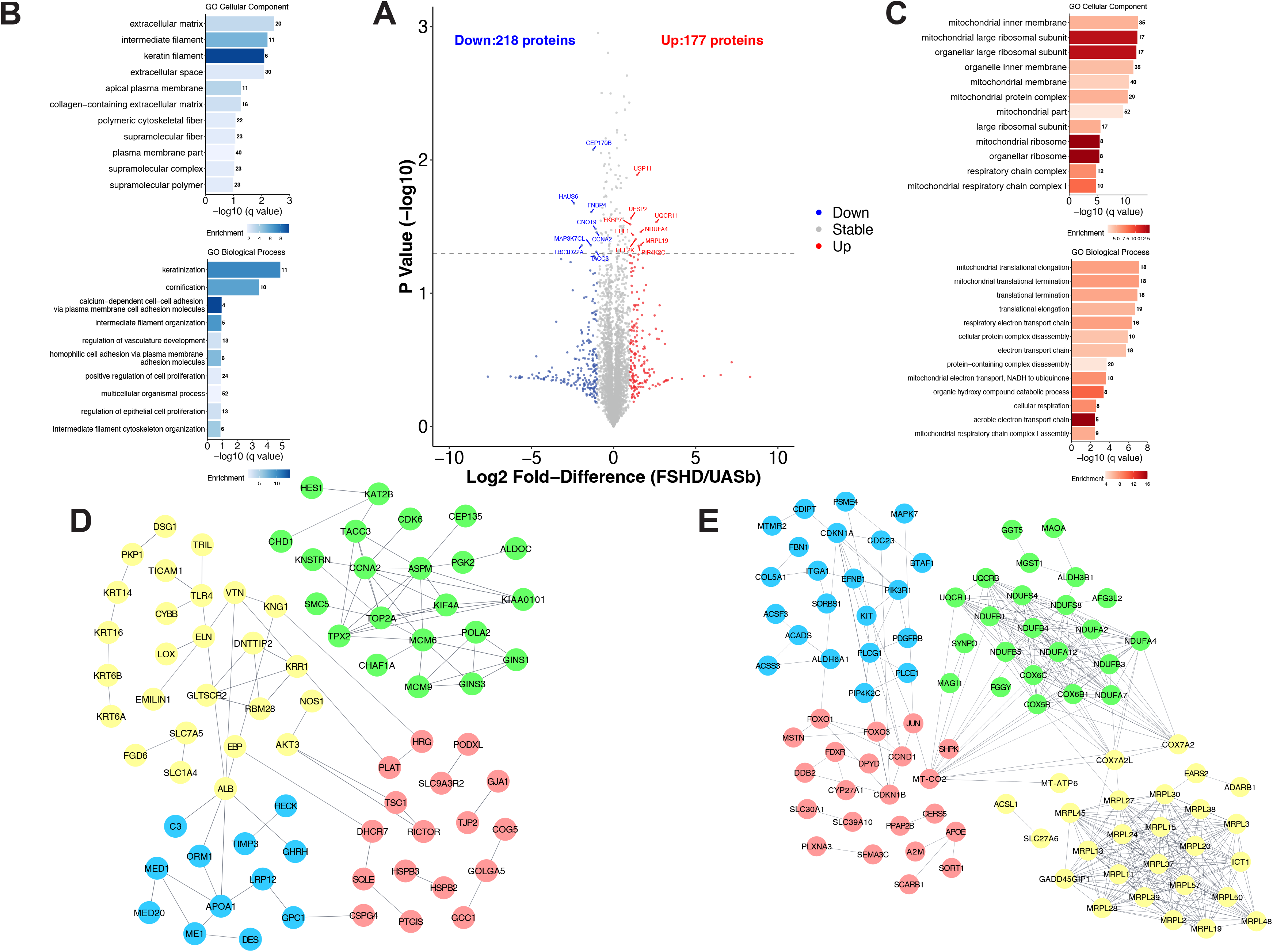
Average protein abundance data. **A)** Volcano plot comparing the Log_2_ Fold-Difference (FSHD/UASb) protein abundance plotted against the -Log_10_ P value (n = 4461). Coloured datapoints represent proteins more abundant (red, log2 Diff. >1), less abundant (blue, log_2_ Diff. <-1), or stable (grey, log_2_ Diff. <1 and >-1) proteins in FSHD myoblasts as compared to UASb. Dashed horizontal line shows a threshold of statistical significance (P < 0.05). Gene ontology (GO) analysis of cellular component and biological process in proteins **B)** depleted (log_2_ Diff. <-1) or **C)** more abundant (log_2_ Diff. >1) in FSHD myoblasts as compared to UASb. GO terms were ranked by -log_10_ (q value) and the number of proteins included into each GO term was shown alongside the bar chart. Each bar chart was filled with the color showing enrichment. STRING protein interaction network in proteins **D)** depleted (log_2_ Diff. <-1) or **E)** more abundant (log_2_ Diff. >1) in FSHD myoblasts as compared to UASb. Full STRING protein interaction network was generated with minimum required interaction score, 0.7. Protein interaction network was clustered using k-means clustering, and each color represents a cluster. Proteins without interaction partners are omitted from visualization.

Gene ontology terms that were significantly enriched amongst proteins depleted (log_2_ Diff. < −1) in FSHD myoblasts, included extracellular matrix and biological processes associated with intermediate filament organization (**Fig. 2B**). Protein interaction networks of those less abundant in FSHD samples formed 4 dominant clusters (**Fig. 2D**), encompassing (i) DNA replication and mitotic cell cycle process (green), (ii) cytoskeleton organization (yellow), (iii) mediator of RNA polymerase II transcription (blue), and (iv) mTOR signaling proteins (red).

Biological processes associated with mitochondrial translation and the respiratory electron transport chain were significantly enriched amongst proteins that were more abundant (log2 Diff. 1) in FSHD myoblasts (**Fig. 2C**). Our analysis encompassed 75 %, 50 %, 80 %, 48 % and 62 % of protein subunits of respiratory chain Complexes I-V, respectively. With the exception of UQCRQ (log_2_ Diff. −0.56) of Complex III and Complex V subunits ATP5PD (log_2_ Diff. −0.16), ATP5ME (log_2_ Diff. −0.59), ATP5F1B (log_2_ Diff. −0.014), all other respiratory chain proteins were more abundant (log_2_ Diff. ranging from 0.1 to 2.6) in FSHD samples.

Specifically, 3 subunits of the respiratory chain complex lV were 2-fold more abundant in FSHD myoblasts, including COX5B, COX7A2, and MT-CO2 though statistical significance was not evident (P>0.05). Sixty-one out of 83 known mitochondrial ribosomal proteins were included in our analysis (73% coverage) and all mitochondrial ribosomal proteins apart from (DAP3, MRPS17, MRPS18B, MRPS25, MRPS28, MRPS35) were more abundant (log_2_ Diff. ranging from 0.016 to 2.5) in FSHD myoblasts as compared to UASb. Specifically, 5 mitochondrial ribosomal proteins were 2-fold more abundant in FSHD myoblasts, including MRPL39, MRPL27, MRPL37, MRPL24, and MRPL3 though statistical significance was not evident (P>0.05). Furthermore, the protein interaction networks derived from proteins that were more abundant in FSHD samples formed 4 dominant clusters (**Fig. 2E**), encompassing (i) mitochondrial ribosomal proteins (yellow), (ii) mitochondrial respiratory chain components (green), (iii) FOXO-mediated transcription (red), and (iv) positive regulation of cyclin-dependent protein kinase activity (blue).

### Differences in protein fractional turnover rate (FTR) between FSHD and UASb myoblasts

Turnover rates were calculated for 2324 proteins that had high-quality peptide mass isotopomer data available at baseline and after 24 h of D_2_O labelling in *n* = 2 FSHD and *n* = 2 UASb samples (**Fig. 3A**). Deuterium incorporation data were unavailable for the 5 proteins that exhibited a significant interaction between group and time or the 11 proteins that exhibited a significant main effect of time therefore the contributions of synthesis and degradation to the changes in protein abundance could not be investigated, unlike our previous study (33). The median turnover rate amongst proteins in FSHD myoblasts (15A: 0.58 %/h, 16A: 0.43 %/h) was not significantly (*P* = 0.37) different from that exhibited by UASb myoblasts (15V: 0.60 %/h, 16U: 0.51 %/h) but the distribution of protein-specific turnover rates in UASb samples was shifted rightward (faster rates of protein turnover) compared to FSHD samples (**Fig. 3C**). One-way analysis of variance (FSHD vs UASb) highlighted 77 proteins that exhibited statistically significant (P<0.05, q>0.99) differences in turnover rate between FSHD and UASb myoblasts (**Fig. 3A**). Thirty-seven of the statistically significant proteins exhibited a >2-fold difference (log_2_ Diff. >1) in turnover, including 11 proteins that had faster rates of turnover in FSHD samples and 26 proteins that had slower rates of turnover compared to UASb samples.

**Figure 3.**
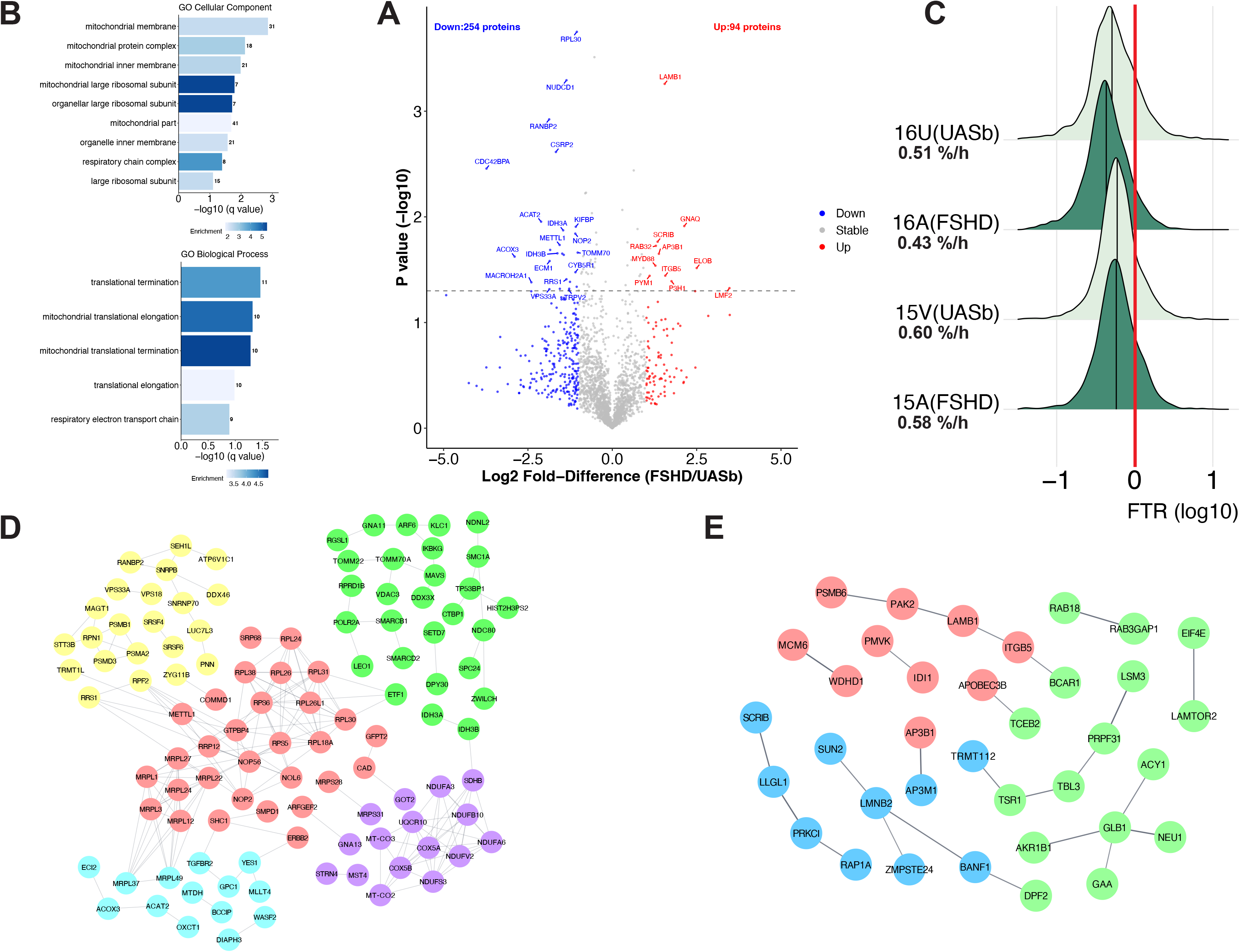
Slower protein turnover rate in FSHD myoblasts as compared to UASb myoblasts. **A)** Volcano plot comparing the Log_2_ fold-change (FSHD/UASb) protein turnover plotted against the -Log_10_ P value (n = 2324). Colored datapoints represent proteins faster (red, log_2_ Diff. 1≥), slower (blue, log_2_ Diff. <-1), or stable (grey, log_2_ Diff. <1 and >-1) turnover proteins in FSHD myoblasts as compared to UASb. Dashed horizontal line shows a threshold of statistical significance (P < 0.05). **B)** Gene ontology (GO) analysis of cellular component and biological process in proteins exhibit slower turnover in FSHD as compared to UASb (log_2_ Diff. <-1). GO terms were ranked by -log_10_ (q value) and the number of proteins included into each GO term was shown alongside the bar chart. Each bar chart was filled with the color showing enrichment. **C)** Density plot of log_10_ transformed fractional protein turnover rate (FTR, %/h) in FSHD and UASb. Vertical line in each density plot indicates the median (the 0.5 quantile) and the red vertical line indicates 1% FTR (0 log_10_). Median values of FTR are shown in bold. No difference in median FTR between FSHD and UASb (one-way ANOVA, P = 0.37). STRING protein interaction network in proteins with **D)** slower turnover rate (log_2_ Diff. <-1) or **E)** faster turnover rate (log_2_ Diff. >1) in FSHD myoblasts as compared to UASb. Full STRING protein interaction network was generated with minimum required interaction score, 0.7. Protein interaction network was clustered using k-means clustering, and each color represents a cluster. Proteins without interaction partners are omitted from visualization.

The Gene ontology terms, including mitochondrial protein complex and mitochondrial large ribosomal subunit and biological processes associated with mitochondrial translational termination and elongation were significantly enriched amongst proteins that exhibited slower (log_2_ Diff. < −1) rates of turnover in FSHD myoblasts. Protein interaction networks that had slower rates of turnover in the FSHD samples formed 5 clusters (**Fig. 3D**), encompassing (i) mitochondrial respiratory chain complex I assembly (purple), (ii) regulation of mRNA processing and proteasome (yellow), (iii) mitochondrial outer membrane translocase complex (green), (iv) ribosomal large subunit assembly and biogenesis (red), and (v) cellular metabolic process (light blue).

No significantly enriched gene ontology terms were detected amongst proteins that exhibited greater (log_2_ Diff. >1) turnover in FSHD myoblasts. Protein interaction networks drawn from proteins that had greater rates of turnover in FSHD compared to UASb samples formed 3 clusters (**Fig. 3E**), which were manually curated as (i) cellular protein localization (green), (ii) extracellular exosome (red), and (iii) tight junction (blue).

### Comparison of protein abundance and turnover data

Differences in protein abundance and turnover data between FSHD and UASb were plotted alongside one another in a scatterplot. Here, the top-left quadrant represents proteins less abundant but greater protein turnover rate in FSHD myoblasts. The top-right quadrant represents proteins more abundant with a greater protein turnover rate in FSHD myoblasts. The bottom-left quadrant represents proteins that were less abundant and exhibited lesser protein turnover in FSHD myoblasts. The bottom-right quadrant represents proteins that were more abundant but exhibit a slower rate of turnover in FSHD myoblasts. More stringent STRING criteria (high confidence (0.7) interaction score, log_2_ Diff. >0.6) were used to highlight core clusters of proteins that share similar patterns across the 2 pairs of patient and sibling samples (**Fig. 4A**). Eleven subunits of mitochondrial Complex I proteins had both abundance and FTR data, and 10 proteins (NDUFA3, NDUFA6, NDUFA8, NDUFA9, NDUFB10, NDUFB11, NDUFS1, NDUFS3, NDUFS6, NDUFV2) were located in the bottom right quadrant (black dots). Two subunits of mitochondrial Complex II proteins had both abundance and FTR data, and SDHB was found in the bottom right quadrant (green dots). Three subunits of mitochondrial Complex III proteins had both abundance and FTR data, and each of these proteins (UQCR10, UQCRC1, UQCRC2) was located in the bottom right quadrant (light blue dots). Six subunits of mitochondrial Complex IV proteins had both abundance and FTR data, and 5 proteins (COX4I1, COX5A, COX5B, MT-CO2, MT-CO3) were located in the bottom right quadrant (purple dots). Notably, mitochondrially encoded gene products, MT-CO2 and MT-CO3 (subunits of mitochondrial Complex IV) were also located in the bottom right quadrant. Abundance and FTR data were measured for 10 subunits of mitochondrial Complex V, and 7 proteins (ATP5F1C, ATP5MF, ATP5MG, ATP5PB, ATP5PD, ATP5PF, ATP5PO) were located in the bottom right quadrant (red dots). Nineteen mitochondrial ribosomal proteins had both abundance and FTR data, and 16 of these (MRPL1, MRPL12, MRPL22, MRPL23, MRPL24, MRPL27, MRPL3, MRPL37, MRPL49, MRPS14, MRPS28, MRPS30, MRPS31, MRPS36, MRPS5, PTCD3) were located in the bottom right quadrant (orange dots). In accordance with the GO analysis of protein abundance and protein turnover data, these proteins are more abundant in FSHD, but have a slower protein turnover rate as compared to UASb. Protein interaction networks (**Fig. 4B**) of proteins that were more abundant (log_2_ Diff. >0.6) and had slower rates of turnover (log_2_ Diff. <-0.6) in FSHD myoblasts formed 2 major clusters, encompassing (i) mitochondrial respiratory chain complex I assembly (yellow) and (ii) mitochondrial translational elongation and termination (red).

**Figure 4.**
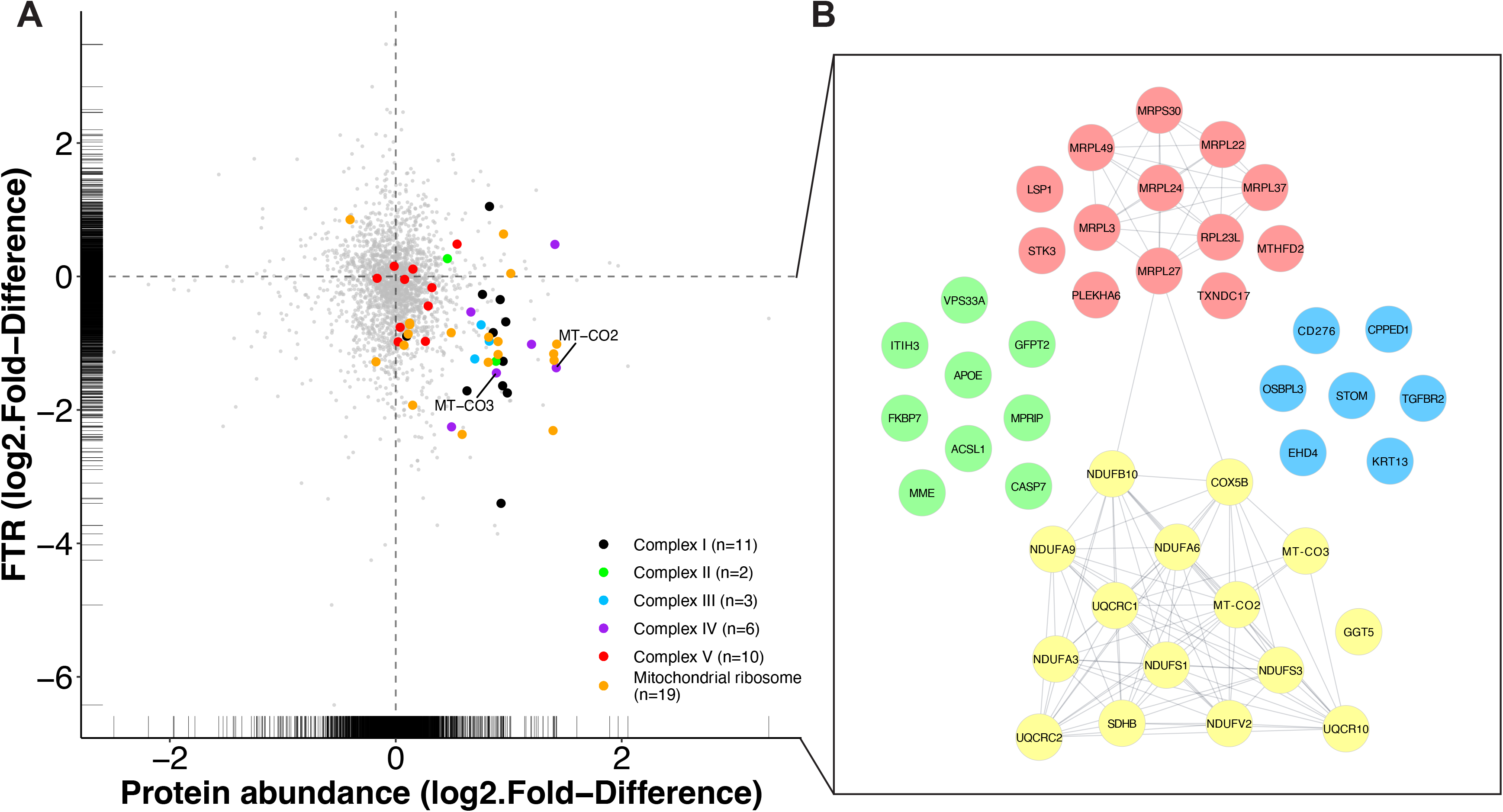
Mitochondrial proteins are more abundant but exhibit slower protein turnover rate in FSHD myoblasts. **A)** Scatter plot comparing the differences in the Log_2_ Fold-Difference (FSHD/UASb) between protein abundance and FTR. Rug plots display distribution of individual data both in X and Y axis. **B)** STRING protein interaction network in proteins more abundant (log_2_ Diff. >0.6) and slower FTR (log_2_ Diff. <-0.6) in FSHD myoblasts as compared to UASb. Full STRING protein interaction network was generated with minimum required interaction score, 0.7. Protein interaction network was clustered using k-means clustering, and each color represents a cluster.

## Discussion

We have used proteomic analysis of stable isotope-labelled patient-derived myoblasts to generate new insight to the potential disease mechanisms of FSHD. Our global proteomic data on the abundance and turnover of individual proteins highlighted that mitochondrial proteins, particularly subunits of the mitochondrial respiratory complexes and mitochondrial ribosomal proteins, were more abundant and had slower rates of turnover in FSHD myoblasts. These findings suggest mitochondrial protein quality might be impaired in FSHD muscle, and FSHD myoblasts may exhibit an accumulation of ‘older’ less viable mitochondrial proteins. Our proteomic dataset also adds new detail to previously known aspects of FSHD muscle pathology, including defects in RNA processing, cell cycle regulation, stress kinase pathways and apoptosis. Our data from two matched pairs of individuals with FSHD and unaffected siblings provide an impetus for further exploration of the role of post-transcriptional processes in pathophysiological mechanisms of FSHD.

Recently, Heher et al. (8) reported the severity of FSHD is positively associated with a lower expression of mitochondrial genes in both skeletal muscle biopsies and myoblasts from individuals affected with FSHD. One of the pairs of FSHD and UASb myoblast samples (16A and 16U) analysed in the current work were also included in the report by Heher et al. (8). Taken together, the differences in gene expression (decreased in FSHD; (8)), protein abundance (increased in FSHD) and protein turnover (decreased in FSHD) of mitochondrial proteins (**Fig. 4**), suggest FSHD myoblasts are characterised by an accumulation of older, potentially less viable, mitochondrial proteins. Most mitochondrial proteins are transcribed from nuclear genes then synthesised in the cytosol and transported into mitochondria, whereas 13 proteins originate from mitochondrial DNA (mtDNA), and contribute subunits to the respiratory chain complexes (43). Our proteomic analysis encompassed approximately half (6 out of 13 proteins) of mtDNA encoded proteins, and each of the mtDNA encoded proteins was more abundant in FSHD myoblasts (**Supplemental Table 1**). Protein turnover data was available for two mitochondrially encoded proteins MT-CO2 (a subunit of Complex lV) and MT-ATP6 (a subunit of Complex V) and each of these proteins were included amongst the cluster of nuclear encoded proteins that were more abundant (>2-fold difference; log_2_ Diff. >1) but had slower rates of turnover in FSHD myoblasts (**Fig. 4 A,B**). Our analysis does not specifically distinguish between mitochondrial proteins that were resident within or outside mitochondria during the study period. Nevertheless, the similarities amongst nuclear-encoded and mitochondrially-encoded proteins may indicate a disease mechanism involving the degradation of proteins within mitochondria rather than an interruption to the transport and degradation of proteins destined for mitochondrial import.

Quadriceps muscle of individuals affected by FSHD exhibits mitochondrial dysfunction and evidence of oxidative stress, including lipofuscin inclusions and protein carbonylation, which correlate with the severity of muscle functional impairments (44). Furthermore, transmission electron microscopy (44) revealed areas of accumulated mitochondrial proteins were associated with myofibrillar disorganization, badly formed mitochondrial cristae, swelling, or separation of the inner and outer membranes in FSHD-affected muscles. Laoudj-Chenivesse et al. (17) reported that proteins involved in mitochondrial oxidative metabolism, such as complex I subunits, NADH dehydrogenase flavoprotein (NDUFV) and NADH-ubiquinone oxidoreductase (NDUFA) are more abundant in both clinically affected and unaffected individuals with FSHD as compared to control subjects. However, mitochondrial disturbances and indicators of oxidative stress were less apparent in clinically affected biceps or deltoid muscle compared to muscles, such as quadriceps, that do not exhibit overt clinical signs of FSHD pathology (17). Previously, the FSHD patient-derived immortalized cells used in this study could not be distinguished from healthy controls based on their response to cellular stresses, including hydrogen peroxide and glutathione depletion (22), unlike other studies that used immortalized cells (5, 20). FSHD myoblasts can repair DNA damage caused by moderate levels of oxidative stress but fail to recover from higher levels or chronic exposure to oxidative stress (45). Using myoblasts from individuals with FSHD and normal control, Winokur et al. (5) demonstrated that FSHD myoblasts are more susceptible to oxidative stress and had lower levels of expression of genes involved in antioxidant processes, such as Glutathione S-transferase theta-2 (GSTT2), Glutathione reductase (GSR), and Heat shock 70 kDa protein 4 (HSPA4). Our proteomic analysis also found that GSTT2 and HSPA4 are less abundant in FSHD myoblasts while GSR was more abundant in FSHD myoblasts (**Supplemental Table 1**), suggesting a heightened oxidative stress and a potential connection to mitochondrial dysfunction.

Apoptosis is one of the affected biological pathways in individuals with FSHD (46, 47) and mitochondria have been shown to contribute to the regulation of apoptosis via mitochondrial derived cytochrome *c* (48). Previous studies suggested that apoptosis can be induced via an increase in mitochondrial ribosome proteins, such as MRPS30 (49) and MRP41 (50). Our proteomic analysis detected a higher abundance of each of these mitochondrial ribosomal proteins in FSHD myoblasts (**Fig. 4B and Supplemental Table 1**). MRPS30 and MRP41 proteins were reported to induce apoptosis via different mechanisms, for example, MRP41 has been reported to induce apoptosis via both p53-dependent (51) and p53-independent (52) pathways. Conversely, Sun et al. (49) demonstrated that overexpression of MRPS30 induces apoptosis while increasing transcription factor c-Jun and by activating of c-Jun N-terminal kinase 1 (JNK1) in mouse fibroblasts. Consistently, our proteomic analysis detected that transcription factor AP-1 (JUN) is enriched in FSHD myoblasts (**Fig. 2E**). Dual specificity protein phosphatase 23 (DUSP23) was also enriched in FSHD myoblasts (**Supplemental Fig. S1 and Supplemental Table 1**) and is known to enhance the activation of JNK, which is a recognized downstream target of DUX4 (53). In support of an activation of apoptosis, several caspase proteins were more enriched in FSHD myoblasts, including caspase-9 (CASP9) and caspase-7 (CASP7) (**Fig. 4B and Supplemental Table 1**), although, the activation status of these proteins cannot be ascertained without knowing their cleavage status. Nevertheless, our data is consistent with recent data (16) linking DUX4 induction with increases in the proportion of Caspase 3/7 positive cells. Several proteins that were uniquely detected in FSHD sample 16A, indicate heightened caspase activity, including protein zyg-11 homolog B (ZYG11B), which is a substrate adaptor subunit in the E3 ubiquitin ligase complex ZYG11B-CUL2-Elongin BC, and plays a role in the clearance of proteolytic fragments generated by caspase cleavage during apoptosis (54). In addition, FOXO1, a transcription factor targeting apoptosis signaling, was particularly more abundant in FSHD sample 16A (FSHD) as compared to 15A (FSHD) (**Supplemental Fig. S1**). These findings suggest sample 16A may exhibit higher caspase activity and a more severe FSHD phenotype than sample 15A.

Antioxidant agents may rescue the impairments in muscle function exhibited by individuals affected with FSHD (55, 56). Antioxidants ameliorate several FSHD-related dysregulated biological pathways, including oxidative DNA damage (6) and apoptosis (57) via a possible suppression of DUX4 transcription. In particular, mitochondrial-targeted antioxidants are more potent than non-targeted agents (8), which suggests respiratory chain dysfunction may be a major contributor to the pathological generation of reactive oxygen species (ROS). Our discovery that subunits of the respiratory chain complexes exhibit a slower rate of turnover in FSHD affected myoblast may point to losses in mitochondrial protein quality control or possibly mitophagy as an underlying mechanism in FSHD. Indeed, Lei et al. (58) report impairments in mitophagy result in an accumulation of dysfunctional mitochondrial excessive mitochondrial ROS generation.

Overexpression of DUX4 in MB135 myoblasts is associated with a 50 % decrease in protein synthesis (14). We also report the median turnover rate of proteins is lesser in FSHD compared to UASb myoblasts (**Fig. 3C**). Our proteomic data reveal protein-specific differences in turnover rate between FSHD and UASb samples (**Fig. 3C**) and discordance between protein turnover and protein abundance data (**Fig. 4A).** The median turnover rate of proteins was less in FSHD sample 16A compared to FSHD sample 15A, and protein mono-ADP-ribosyltransferase PARP12 (PARP12) was specifically detected in 16A (FSHD) (**Supplemental Fig. S1**). PARP12, a member of a large family of ADP-ribosyl transferases, can be recruited to stress-granules (i.e., known sites of mRNA translation arrest) and block mRNA translation (59), which may contribute to the greater impairment of protein synthesis FSHD sample 16A. In agreement with Jagannathan et al. (14), we found proteins involved in the negative regulation of protein synthesis were more abundant in FSHD myoblasts, including eukaryotic elongation factor 2 kinase (EEF2K) (**Fig. 2A**), which inhibits translation elongation. Jagannathan et al. (14) reports the phosphorylation of eIF2a is increased in a time dependent manner after DUX4 induction (i.e., indication of translation inhibition) with a concomitant decrease in [^35^S]-Methionine incorporation. In our data, other proteins associated with the positive regulation of protein synthesis were less abundant in FSHD myoblasts, including large neutral amino acids transporter small subunit 1 (SLC7A5), RAC-gamma serine/threonine-protein kinase (AKT3), Hamartin (TSC1), and Rapamycin-insensitive companion of mTOR (RICTOR) (**Fig. 2D and Supplemental Table 1**). However, the kinase activity of these proteins involved in mTOR signaling pathway remains to be investigated.

FSHD is associated with a progressive decline in muscle mass and our proteomic analysis detected proteins associated with negative regulation of skeletal muscle growth, including Growth/differentiation factor 8 (MSTN) and Tensin-2 (TNS2), which were more abundant in FSHD myoblasts (**Fig. 2E and Supplemental Table 1**). Moreover, the GO biological process-terms depleted in FSHD myoblasts included positive regulation of cell proliferation (**Fig. 2B**), based on the finding that Cyclin-dependent kinase 6 (CDK6), Cyclin-A2 (CCNA2), Cyclin-dependent kinase 13 (CDK13), Cyclin-dependent kinase 9 (CDK9) were decreased (**Fig. 2D and Supplemental Table 1**). Conversely, proteins associated with a negative regulation of cell cycle, such as Cyclin-dependent kinase inhibitor 1B (CDKN1B), Cyclin-dependent kinase inhibitor 1 (CDKN1A), Cyclin-dependent kinase 4 inhibitor (CDKN2C), Cyclin-dependent kinase inhibitor 1C (CDKN1C), Cyclin-dependent kinase 4 inhibitor B (CDKN2B) were increased (**Fig. 2E and Supplemental Table 1**). Cyclin-dependent kinase (Cdk) is activated by association with cyclin subunits and phosphorylation of the Cdk subunit by the Cdk-activating kinase. Thus, Cdk activity and the profile of cell cycle phases warrant further investigation in FSHD and UASb myoblasts.

Human myogenic cells expressing DUX4-FL exhibit insoluble ubiquitylated proteins (46), which suggests the ubiquitin proteasome system may be impaired in FSHD. Furthermore, proteins involved in the ubiquitin proteasome system are increased at the transcriptional and protein level (14, 60, 61) after induction of DUX4. In our data, several ubiquitin E2 conjugating enzymes (UBE2T, UBE2D2, UBE2Q1), ubiquitin E3 ligases (UBR3, UBR7, XIAP, TRIM32, RNF149, RNF213), deubiquitylating enzymes (USP11, USP8, USP22, OTULIN, UCHL1), as well as proteasome activator complex subunit protein (PSME4) were each more abundant in FSHD myoblasts (**Fig. 2 A,E and Supplemental Table 1**) but, nevertheless, protein turnover rates were generally lower. In addition, our proteomic analysis detected that Forkhead box protein O1 (FOXO1) and Forkhead box protein O3 (FOXO3) were more abundant in FSHD myoblasts (**Fig. 2E and Supplemental Table 1**). These FOXO1 and FOXO3 transcription factors (62) are master regulators of various ubiquitin E3 ligase proteins (63, 64) and autophagy lysosome (65, 66). Moreover, we found that proteins involved in autophagy lysosome system (TM9SF1) and protein quality control (HSPA1L, HSPA14, SGTA) are also more abundant in FSHD myoblasts (**Supplemental Table 1**). Thus, our data potentially indicate a failed attempt to maintain or alter muscle proteostasis in FSHD myoblasts.

Our proteomic analysis did not detect previously suggested FSHD or DUX4 candidate genes highlighted in the biomarker list reported in Yao et al. (67). This is unsurprising given that DUX4 is transiently expressed and challenging to detect in FSHD patient samples. Moreover, we studied FSHD myoblasts in growth media, conditions which are not expected to be associated with high levels of DUX4 expression (30). Nevertheless, our proteomic data align well with the wider literature on the molecular mechanisms of FSHD. Notably, when DUX4 is artificially expressed in human myoblasts, only 8 of 25 candidate genes (67) were detected at the protein level (14). Moreover, almost half (6 of 13) of the FSHD patient samples reported by Yao et al. (67) did not show enrichment of the list of DUX4 target genes. The mitochondrial ADP:ATP carrier, ANT1 (also known as SLC25A4), resides in the 4q35 region alongside DUX4 (17) and has also been a focus of interest in FSHD research. We did detect ANT1 but did not find significant differences in either the abundance or turnover rate of ANT1 between FSHD and UASb myoblasts. Our ANT1 data agree with Klooster et al. (68) but contrast with Gabellini et al. (69) that demonstrated that ANT1 was more abundant in the muscle of both in clinically affected and unaffected individuals with FSHD compared to healthy control subjects.

## Conclusions

Dynamic proteome profiling has offered new insight to the disease mechanisms of FSHD that are underpinned by post-transcriptional processes. We discovered that FSHD myoblasts exhibit a greater abundance but slower turnover rate of mitochondrial respiratory complex subunits and mitochondrial ribosomal proteins, which may indicate an accumulation of ‘older’ less viable mitochondrial proteins. Our data provide a substantial hypothesis-generating resource for the FSHD community and highlight the importance of post-transcriptional processes and protein turnover in FSHD pathology.

**Figure 5.**
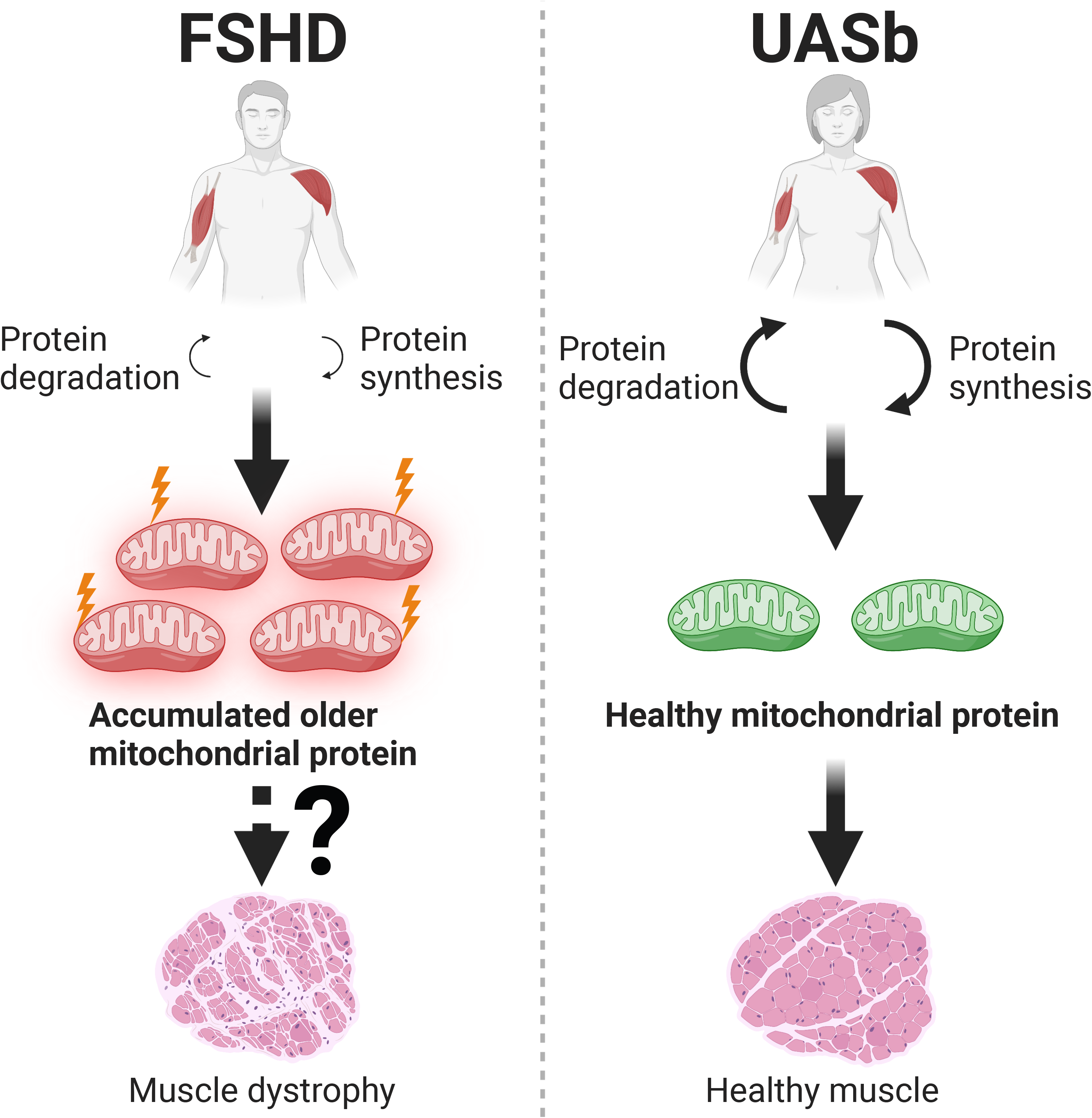
Accumulation of “older” less viable mitochondrial proteins may contribute to the pathophysiology of FSHD. FSHD myoblasts exhibit a greater protein abundance but slower turnover rate of mitochondrial respiratory complex subunits and mitochondrial ribosomal subunits, which may indicate an accumulation of ‘older’ less viable mitochondrial proteins compared to UASb myoblasts. This may contribute to the reduced respiratory function specifically observed in Complex I as recently shown by Heher et al. (8) in DUX4 expressing iDUX4 myoblasts. Impaired mitochondrial function is proposed as one of the major pathophysiological mechanisms of FSHD.

## Abbreviations

D_2_O: deuterium oxide
FSHD: facioscapulohumeral muscular dystrophy
FASP: Filter-Aided Sample Preparation
DUX4: double homeobox 4
FTR: fractional turnover rate
BSA: bovine serum albumin
DTT: dithiothreitol
Ambic: ammonium hydrogen bicarbonate
TFA: trifluoracetic acid
GO: gene ontology
UASb: unaffected siblings
DUX4-FL: full-length isoform of DUX4
mtDNA: mitochondrial DNA
ROS: reactive oxygen species

## Data availability

The mass spectrometry proteomics data generated in this study have been deposited to the ProteomeXchange Consortium via the PRIDE (70) under the identifier PXD038818 and 10.6019/PXD038818.

## Supplemental data

This article contains supplemental data.

## Funding and additional information

Muscular Dystrophy Association – Strength, Science, and Stories of Inspiration Fellowship (AJB), Muscular Dystrophy UK (YN, Y-WC and JGB), FSHD Canada (Y-WC), FSHD Society Grant Award (AJB), and American Physical Therapy Association New Investigator Fellowship Training Initiative (AJB).

## CRediT author statement

Conceptualization (JGB, Y-WC), Data Curation (JGB), Formal Analysis (YN, AJB, CAS, JGB), Funding Acquisition (Y-WC), Investigation (YN, AJB, CAS), Methodology (AJB, Y-WC, JGB), Project Administration (Y-WC), Resources (Y-WC, JGB), Software (JGB), Supervision (JGB, Y-WC), Visualization (YN, CAS), Writing – Original Draft Preparation (YN, JGB), Writing – Review & Editing (YN, AJB, CAS, JGB, Y-WC)

## Conflict of interest

Authors declare no competing interests.

## Notes

### Competing Interest Statement

The authors have declared no competing interest.

