## Supplemental Figure 1 for "Facioscapulohumeral muscular dystrophy is associated with altered myoblast proteome dynamics"

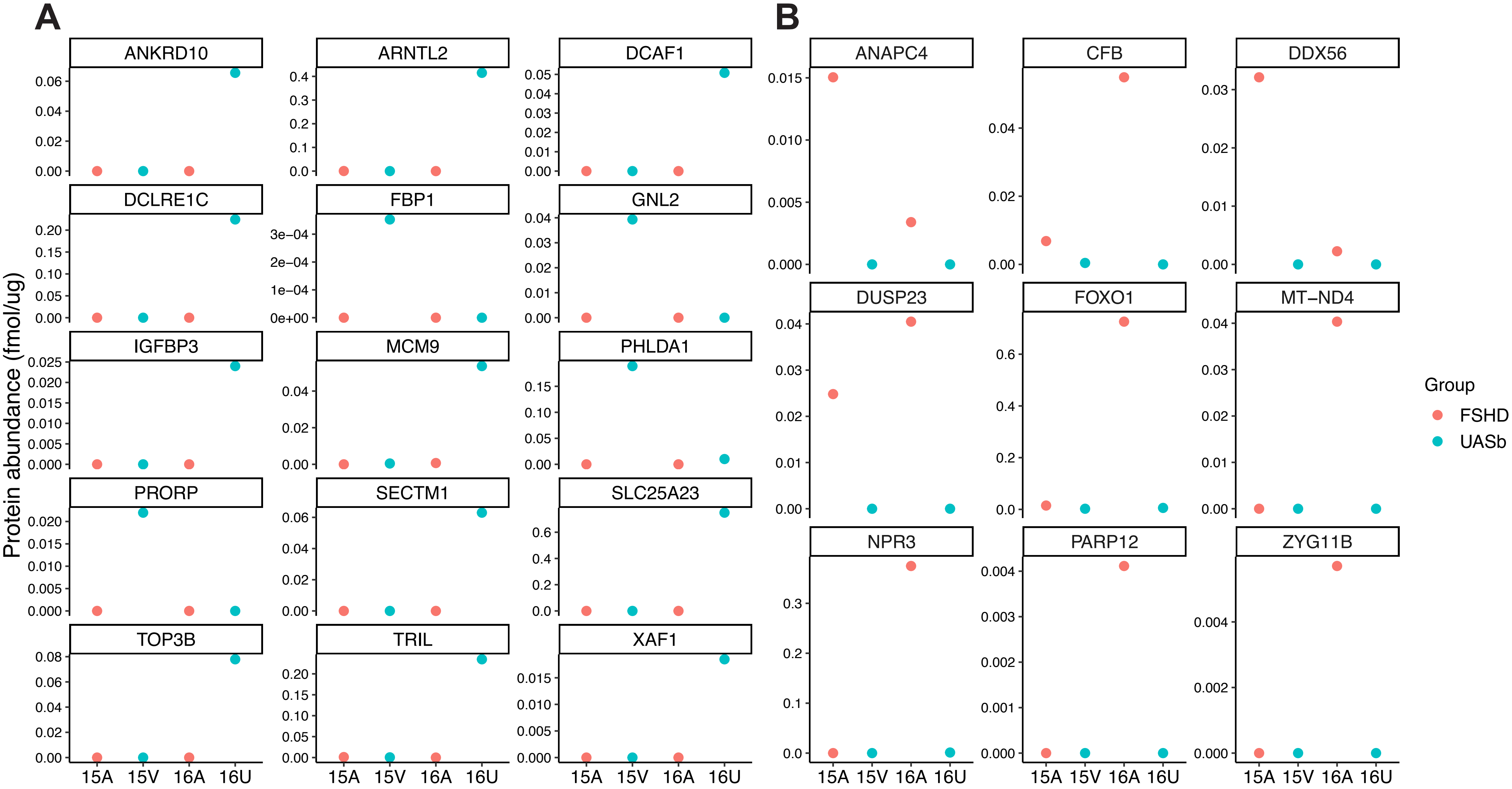


**Fig. S1**: *Family and group-specific characteristic of protein abundance*. Proteins specifically detected in A) UASb or B) FSHD.
